# Memory consolidation during sleep involves context reinstatement in humans

**DOI:** 10.1101/2022.03.28.486140

**Authors:** Eitan Schechtman, Julia Heilberg, Ken A. Paller

**Author notes:** **Corresponding author and lead contact:** Eitan Schechtman.

## Abstract

New memories aren’t quarantined from each other when first encoded; rather, they are interlinked with memories that were encoded in temporal proximity or share semantic features. By selectively biasing memory processing during sleep, here we test whether context influences sleep-consolidation. Participants first formed 18 idiosyncratic narratives, each linking four objects together. Before sleep,they also memorized an on-screen position for each object. During sleep, 12 object-specific sounds were unobtrusively presented, thereby cuing the corresponding spatial memories and impacting spatial recall as a function of initial memory strength. As hypothesized, we find that recall for non-cued objects contextually linked with cued objects also changed. Post-cue electrophysiological responses suggest that activity in the sigma band supports context reinstatement and predicts context-related memory benefits. Concurrently, context-specific electrophysiological activity patterns emerge during sleep. We conclude that reactivation of individual memories during sleep evokes reinstatement of their context, thereby impacting consolidation of associated knowledge.

## Introduction

Individual memories are supported by an intricate network of interconnections—not independent and detached from other memories. These connections are central to the organization of memories in the brain and impact subsequent retrieval. The term “context” has been used to describe the elements that surround a core memory, which share some features with it such as time, space, or semantic-relatedness^1,2^. Memories that are formed within temporal proximity of others are said to share a temporal context, whereas memories that share semantic relatedness are said to share a semantic context or be semantically clustered. Both types of contexts impact subsequent retrieval^2,3,4,5,6^. When a specific memory learned in some context is retrieved (e.g., the decorations for a recent party), other contextually related memories may effortlessly come to mind as well (e.g., the guests attending the party). On the neural level, this process, termed contextual reinstatement, is manifested by increased similarity between the observed neural patterns during encoding and retrieval^7,8^. In this study, we explored contextual reinstatement when memories were reactivated during sleep, as well as the consequences of reinstatement on retrieval.

Whereas the role of context at encoding and retrieval has been repeatedly demonstrated, its role in the intermediate period has not been systematically explored. During these offline periods, including sleep, memory traces are cemented in cortical networks through a set of processes collectively termed consolidation^9,10^. The consolidation of declarative memories (i.e., explicit memories for facts and autobiographical events) is thought to primarily occur during non-rapid-eye-movement (NREM) sleep, which consists of stages N2 and N3 of sleep. Consolidation is thought to rely on memory reactivation, which, like contextual reinstatement, involves the selective activation of memory-specific neural circuits^11,12^. The extent to which contextually related memories are reinstated over the course of consolidation during sleep remains unclear. Recently, it has been hypothesized that the benefits of NREM to memory stem from it being a state devoid of context, thus preventing the damaging effects of contextual interference^13^ (but see^14^).

To explore context reinstatement during sleep, we biased memory processing during sleep using unobtrusive stimuli, a technique termed targeted memory reactivation (TMR)^15^. TMR has been used to improve different forms of memory, including declarative and nondeclarative^16^. Using this causal manipulation, we demonstrated that consolidation was not limited to memories that were directly targeted, but extended to other memories that were contextually bound to them. These results suggest that context plays a role in the process of memory consolidation during sleep. By examining electrophysiological waveforms following stimulus presentation during sleep, we demonstrated that power in the sigma range (15-20 Hz)—which may reflect the activity of sleep spindles linked with memory consolidation^17,18,19^—reflects the process of contextual reinstatement during sleep and predicts subsequent performance on a memory task.

## Results

Participants (N=29) engaged in a single-session experiment that included an afternoon nap (Figure 1a). They first invented idiosyncratic stories involving a place (e.g., a movie theater) and four objects (Figure 1b). Next, they encoded the positions of objects on a 2-d grid (Figure 1c). To keep the encoding context (i.e., the story) salient, this training also involved answering story-specific questions that required contextual reinstatement. Spatial recall was then tested for all object positions. The average positioning error, calculated as the Euclidean distance between the true position and placed position, was 76.7±1.9 (mean ± SEM) pixels (see STAR Methods). Next, participants were allowed to nap for up to 90 minutes. During NREM sleep, participants were exposed to sounds that had been associated with some of the bjects (Supplementary Table 1). After sleep, spatial recall was once again tested for all object positions. Finally, as a manipulation check, participants engaged in a cued recall test, in which they were asked to list the objects linked with each scene. On average, participants recalled 3.79±0.05 objects out of the 4 associated with each scene, with no effect of cuing on recall (*p* = 0.91).

**Figure 1:**
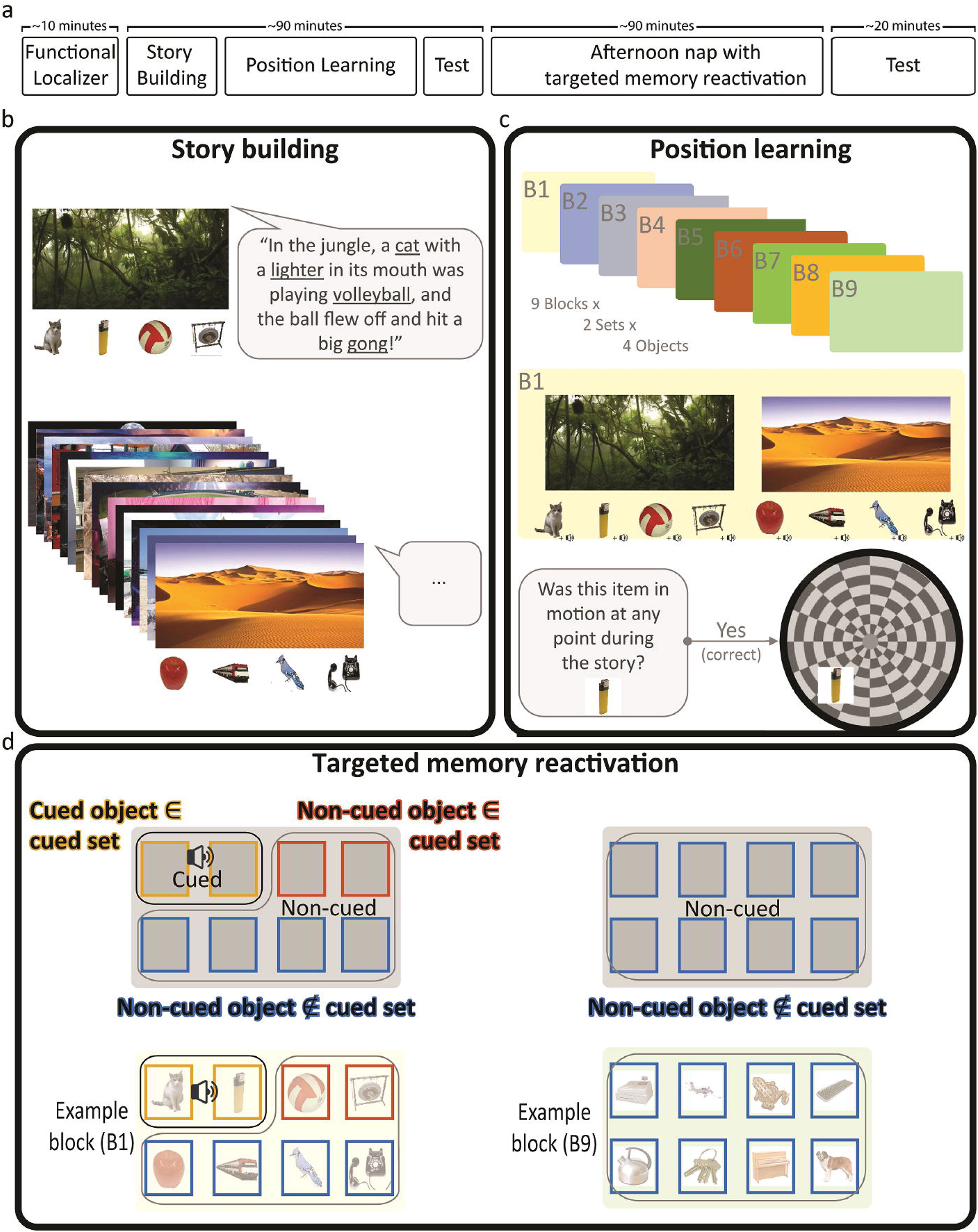
Experimental design. (a) Participants underwent tasks before and after a 90-minute afternoon nap. (b) Participants constructed a story including the four objects and the locale pictured. The four objects, linked together by the story, formed a contextually bound set. Participants constructed 18 such stories. (c) Participants learned the on-screen positions of the eight objects in conjunction with answering questions about their stories. Each of nine blocks featured two contextually bound sets. (d) After a memory test for object positions, participants had a 90-minute nap opportunity. During their sleep, sounds associated with 12 objects were unobtrusively presented. These objects were systematically selected from six of the nine blocks. In these six blocks, two objects from the same set were cued. The critical conditions were thus: cued objects, non-cued objects from a set with cued objects (i.e., a cued set), and non-cued objects that do not belong to a cued set (either from a cued block or a non-cued block). The panel shows an outline of the two types of blocks (top) and example blocks for each (bottom).

### Reactivation during sleep impacted performance for contextually bound memories

We hypothesized that biasing reactivation for certain memories would impact other memories that belonged to the same contextually bound set. We first divided objects for each participant into three groups (Figure 1d): (1) objects that were directly reactivated using sounds during sleep (cued objects ∈ cued set); (2) objects that were not directly reactivated, but were contextually linked with objects belonging to the previous group (non-cued objects ∈ cued set); (3) objects that were neither directly reactivated nor shared an encoding context with those that were (non-cued objects ∉ cued set) (Figure 1d). The three groups did not differ in terms of pre-sleep accuracy errors (*F*(2,1666)=1.29, *p*=0.28; average error were 71.53±3.26, 75.66±3.57, 77.89±2.05, respectively). After sleep, the average errors for the three groups were 80.55±3.26, 82.47±4.31, 81.75±1.95, respectively.

We hypothesized that the second group would show less forgetting during sleep relative to the third group. We first submitted our results to a repeated-measures ANOVA across participants with Condition (i.e., the three groups of objects) and Time (pre- and post-sleep) as factors. We found a main effect of Time (*F*(1,56)=7.51, *p*<0.05), indicating that errors increased across sleep. The effects of Condition and the interaction between Condition and Time were not significant (*F*(2,56)=0.94, *p*=0.34, *F*(2,56)=0.45, *p*=0.51, respectively). We verified these results by running three additional repeated-measures ANOVAs contrasting all three possible group pairings (e.g., cued objects ∈ cued set vs. non-cued objects ∉ cued set). None of these analyses yielded significant effects of Condition or significant interactions (all *p*s>0.14). Diverging from our prediction and from previous studies using targeted memory reactivation, these results suggest that reactivation during sleep did not impact memory. In fact, forgetting rates were numerically higher for cued objects relative to non-cued ones, although this difference was not significant. Since reactivation did not improve memory uniformly, we next turned our attention to its effects on memories as a function of their initial encoding strength.

Building on prior studies showing that sleep’s impact on memory is modulated by pre-sleep memory strength^20,21,22,23^, we hypothesized that the effect of cuing in our task may depend on the initial memory strength as reflected by pre-sleep errors. We fitted a mixed linear model predicting how post-sleep positioning errors would be modulated by Condition, while accounting for pre-sleep positioning errors (Figure 2a, b). This approach allowed us to consider whether reactivation affected memory differentially based on accuracy levels before sleep. We thus focused on the interaction between pre-sleep error and condition, which we term encoding-strength-dependent forgetting (ESDF). Higher ESDF values indicate more forgetting for objects with weaker encoding strength and less forgetting for stronger memories.

**Figure 2:**
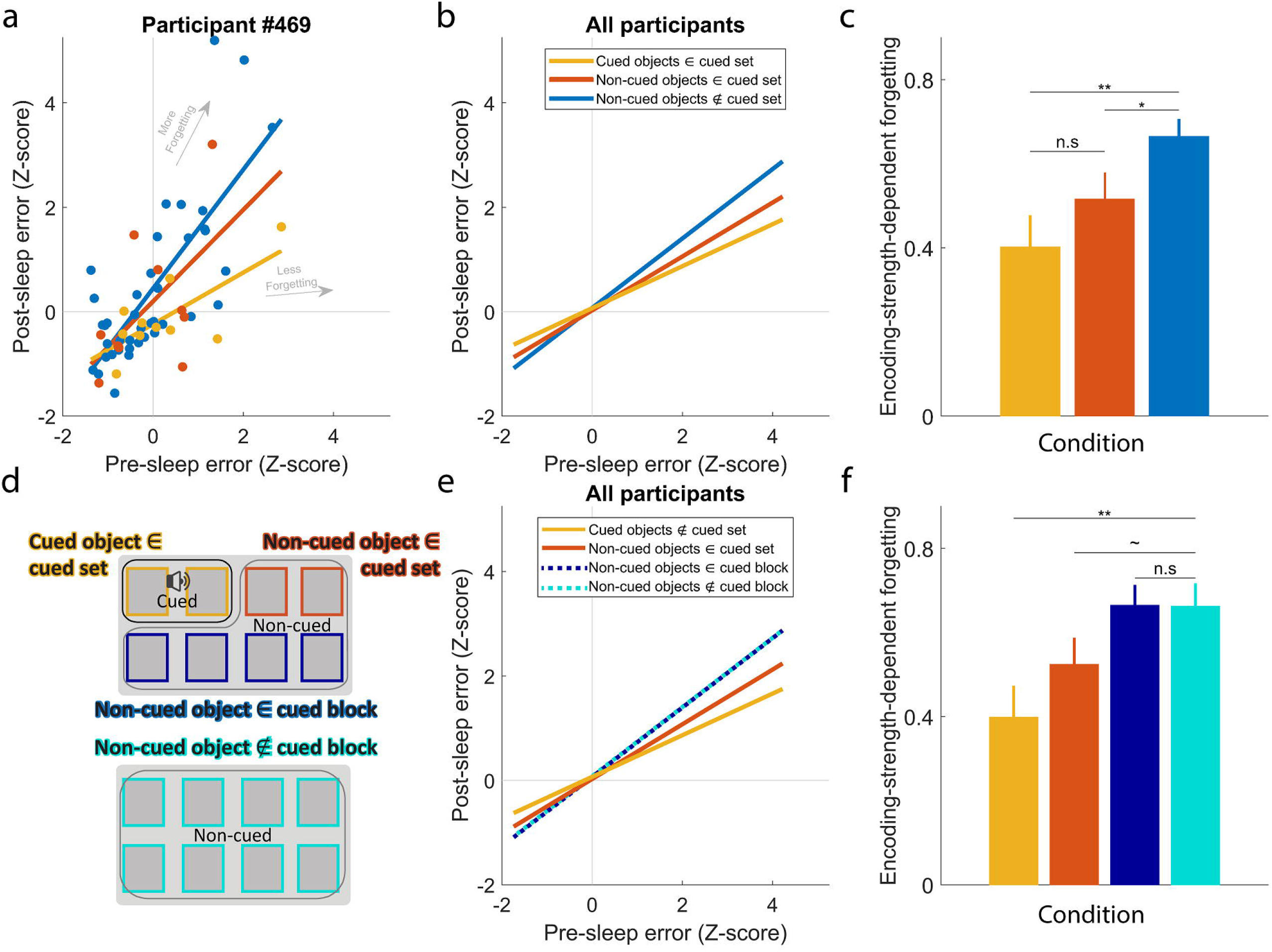
Targeted reactivation impacted memory for reactivated memories and for memories bound by semantic context – but not temporal context. (a) Data for a single participant. Each dot represents error rates for objects positioned by the participant before and after sleep. Colors signify the condition for each object: cued objects ∈ cued set (yellow), non-cued objects ∈ cued set (red), non-cued objects ∉ cued set (blue). Steeper slopes signify more encoding-strength-dependent forgetting between the pre- and post-sleep tests (e.g., memory for the weakly encoded objects gets worse). (b) Forgetting curves across all participants. Slopes and intersects were estimated using a mixed linear model. (c) Statistical comparison of encoding-strength-dependent forgetting (ESDF; slopes in Figure 2b) across conditions. Higher values reflect more forgetting for weaker memories and less forgetting for stronger memories. (d) The same dataset was submitted to another analysis, which distinguishes between two groups of objects: [(non-cued objects ∉ cued set) ∈ cued block] and (non-cued objects ∉ cued block). For brevity, the former group is designated as (non-cued objects ∈ cued block). Conventions follow Figure 1d. (e) ESDF values across all participants for the four conditions. The purple and turquoise lines largely overlap but reflect data for different objects. (f) Statistical comparison of ESDF values across conditions. Error bars signify standard errors of the mean and p-values reflect contrasts between parameters produced using mixed linear models. **-*p*<0.01; *-*p*<0.05; ∼-*p*<0.1; n.s-*p*>0.1.

Normalized pre-sleep errors were positively correlated with post-sleep errors (*F*(1,1537)=258.8, *p*<0.001), indicating that memories that were well-remember before sleep remained relatively well-remembered after sleep (and vice versa). Mirroring the results of the repeated-measures ANOVA, we did not find a main effect of Condition (*F*(2,1537)=0.28, *p*=0.75), indicating that cuing did not reduce post-sleep errors uniformly. However, we found a significant interaction between Condition and pre-sleep errors (*F*(2,1537)=7.1, *p*<0.001), indicating that cuing differentially impacted errors as a function of pre-sleep errors, with greater benefits for weakly encoded memories and fewer benefits for strongly encoded ones. This interaction reflects differences between conditions in encoding-strength-dependent forgetting. Higher ESDF values reflect more forgetting for objects with higher pre-sleep errors (i.e., steeper slopes between pre- and post-sleep errors as shown in Figure 2a, b). We compared encoding-strength-dependent forgetting values for the three conditions to unpack the interaction effect (Figure 2c). Directly reactivated memories (i.e., cued objects ∈ cued set) showed smaller ESDF values relative to memories that were neither reactivated nor contextually bound to reactivated memories (i.e., non-cued objects ∉ cued set; *t*(1537)=-3.1, *p*<0.01). Crucially, non-cued objects that were contextually bound with cued objects (i.e., non-cued objects ∈ cued set) showed smaller ESDF values relative to memories that were neither reactivated nor contextually bound to reactivated memories (i.e., non-cued objects ∉ cued set; *t*(1537)=-2.2, *p*<0.05). ESDF values for cued and non-cued memories within the same contexts were not significantly different (i.e., cued objects ∈ cued set vs non-cued objects ∈ cued set ; *t*(1537)=-1.07, *p*=0.29). Comparable results were obtained when considering the non-normalized accuracy errors, measured in pixels (Supplementary Figure 1). Taken together, these results indicate that the effects of memory reactivation during sleep extend beyond targeted memories, impacting other memories that were contextually linked with them.

ESDF values may reflect forgetting rates for weaker memories (for which lower ESDF values reflect less forgetting), stronger memories (for which lower ESDF values reflect more forgetting), or a combination of both. To explore these effects separately, we used a median split (calculated separately for each participant based on their data distribution in T1) to examine the effects of reactivation on memories with higher-than-median and lower-than-median error rates. A linear mixed model incorporating median in lieu of T1-error showed a significant interaction between Condition and Median-split (*F*(2,1537)=4.53, *p*<0.05), mirroring the interaction between Condition and T1-error reported in the previous paragraph. We then analyzed data for the higher-than-median and lower-than-median errors separately to ask whether the interaction is driven by a main effect of condition on weaker or stronger memories, respectively. However, neither half revealed a significant effect of Condition (*p*>0.2). There was thus no evidence that ESDF values were driven by an effect isolated to weaker or stronger memories, leaving open the possibility that a composite from both was operative.

Using the same dataset, we next examined the differential roles of semantic context (i.e., conceptual links between objects, operationalized by the idiosyncratic story for each set) and temporal context (i.e., links between objects learned within temporal proximity, operationalized by our block design; Figure 1c). Temporal encoding context has been shown to reinstate during wake, thereby impacting retrieval^8,24,25^, and we hypothesized that it would similarly be reinstated and impact reactivation during sleep. We divided our data into four groups (Figure 2d). The first two were identical to those used for the previous analysis (cued objects ∈ cued set; non-cued objects ∈ cued set). Another group included objects that were not linked with a cued set, but were learned within the same block as cued objects [(non-cued objects ∉ cued set) ∈ cued block]. The final group included objects that were neither learned in the same block nor linked with the same set as the cued objects (non-cued objects ∉ cued block). Objects in the former group – but not the latter group – were learned within temporal proximity (i.e., within the same block) of cued objects. We postulated that these temporally segmented blocks would effectively create segregated temporal contexts that would drive behavior.

The four groups did not differ in terms of pre-sleep accuracy errors (*F*(3,1665)=0.91, *p*=0.43; average errors were 71.53±3.26, 75.66±3.57, 77.64±2.82, 78.12±2.19, respectively). After sleep, the average errors for the four groups were 80.55±3.26, 82.47±4.31, 80.89±2.89, 82.46±2.21, respectively. Like before, we first submitted our results to a repeated-measures ANOVA across participants with Condition (i.e., the four groups of objects) and Time (pre- and post-sleep) as factors. We found a main effect of Time (*F*(1,84)=9.16, *p*<0.01), indicating that errors increased across sleep. The effects of Condition and the interaction between Condition and Time were not significant (*F*(3,84)=0.76, *p*=0.39, *F*(3,84)=0.51, *p*=0.48, respectively). Using a linear model to investigate encoding-strength-dependent effects, we found main effects of pre-sleep errors (*F*(1,1535)=150.2, *p*<0.001) but not condition (*F*(3,1535)=0.2, *p*=0.9), as well as a significant interaction (*F*(3,1535)=4.6, *p*<0.01; Figure 2e). Contrasting the ESDF values for the three groups revealed no significant difference between the [(non-cued objects ∉ cued set) ∈ cued block] and (non-cued objects ∉ cued block) groups, suggesting that temporal context, as operationalized by learning block, had no effect on performance (*t*(1535)=0.03, *p*=0.98; Figure 2f). We found significantly higher ESDF values for the (cued objects ∈ cued set) group relative to the (non-cued objects ∉ cued block) group (*t*(1535)=-2.89, *p*<0.01), and a trend toward higher ESDF values for the (non-cued objects ∈ cued set) group relative to the (non-cued objects ∉ cued block) group (*t*(1535)=-1.83, *p*=0.07).

Taken together, results suggest that semantic context is reinstated during sleep and guides memory consolidation. Temporal context did not show any effects on consolidation, but it is possible that such effects may have been missed due to shortcomings of our operationalization of temporal context in this paradigm.

### Sigma oscillations reflect contextual reinstatement, predicting changes in performance

Next, we explored the role of sleep EEG oscillations in the process of contextual reinstatement. First, we calculated the time-locked time-frequency response to sounds presented during sleep for all task-related sounds (Figure 3a). Across participants, we identified two clusters in the time-frequency representation following sound onset (*p* < 0.01, corrected): one cluster at lower frequencies (<10 Hz) peaking around 0.5 s after sound onset, and another between 15-20 Hz, which involved two components peaking before and after the 1 s mark after sound onset (Figure 3b, left). Cluster 1 (Figure 3b, right), consisting of frequencies in the delta and theta range, putatively reflects activity related to slow oscillations and K-complexes, which are typical of NREM sleep. Cluster 2, consisting of frequencies in the sigma range, may reflect the occurrence of sleep spindles, a sleep-specific waveform which has been linked to consolidation^17,18^. However, previous research has shown that spindles typically commence approximately 1 s after sound onset^11,19,26^. Together with the conjoined pattern of the observed cluster, this may suggest that only the late component of the cluster (cluster 2B) truly reflects spindle activity, whereas the early component (cluster 2A) reflects some high-frequency component of the K-complex^27^. Therefore, we conducted further analyses on both the full high-frequency cluster and its two components.

**Figure 3:**
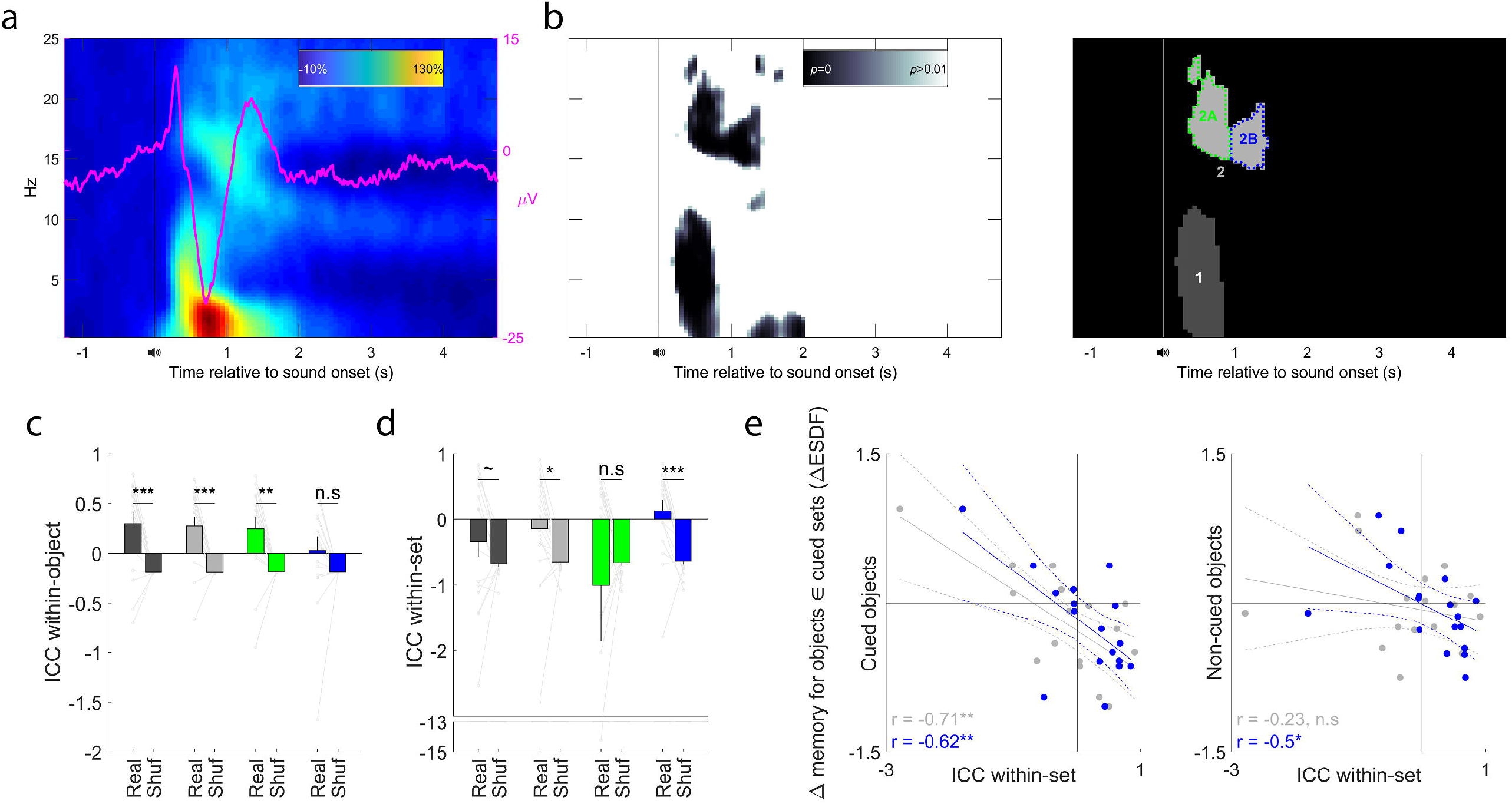
Post-cue spectral power is driven by context and predicts contextually driven changes in performance. (a) Time-frequency representation of EEG activity from electrode Cz over the time period before and after cue onset during sleep. The pink line represents the average event-related response. (b) Left – map of the across-participant p-values for changes in spectral power. Right – identified clusters. (c) The similarity of induced power changes between different repetitions of the same sound was quantified using intraclass correlation coefficients for each cluster. Bar colors correspond to cluster outline colors in panel b (i.e., clusters 1, 2, 2A, and 2B from left to right). Results were evaluated using a permutation test. Real – values obtained for non-shuffled data. Shuf – values obtained for shuffled data. Gray lines represent individual participant data (p-values obtained from paired t-tests between the shuffled and real values). (d) The similarity of induced power changes between different sounds belonging to the same contextually bound set. Designations follow those in panel c. (e) Correlations between intraclass correlation coefficients and condition-specific changes in encoding-strength-dependent forgetting values across participants. Left – for cued objects ∈ cued sets. Right – for non-cued objects ∈ cued sets. P-values reflect the significance level of the Pearson correlation coefficients. Error bars and dashed lines signify standard errors of the mean. ***-*p* < 0.001; **-*p*<0.01; *-*p*<0.05; ∼-*p*<0.1; n.s-*p*>0.1.

Throughout NREM sleep, sounds were often presented multiple times (Supplementary Table 1). We hypothesized that waveforms resulting from repeated presentations of the same sound would be more similar one to the other than those resulting from different sounds. To test this idea, we first quantified power in each cluster on a trial-by-trial basis. Per trial, the average power within the cluster in time-frequency space was averaged, to create a single scalar for each cluster in each trial. These values obtained for each cluster in each participant were submitted to an intraclass correlation coefficient (ICC) analysis (Figure 3c). ICC is a metric that quantifies within-vs between-group correlations. In this case, higher ICC values would indicate that the response was more consistent between repetitions of the same sounds (e.g., all *cat* sound presentations) than between different sounds (e.g., *cat* vs *lighter* sound). For most clusters, results indeed indicated that coefficients were higher than those expected by chance, as assessed using a permutation test (cluster 1, *p*<0.001; cluster 2, *p*<0.01; cluster 2A, *p*<0.01; cluster 2B, *p*=0.15 ; Figure 3b). These correlations may be the result of the memory content related to a sound being reactivated similarly across trials, but a more parsimonious explanation is that they stem from the acoustic properties of the presented sounds creating similar electrophysiological responses.

We used a similar approach to test whether specific waveforms reflected contextual relationships between objects within the same set. We hypothesized that responses would be more similar across sounds if these sounds were linked within the same contextually bound set. For example, if a *cat* object and a *lighter* object shared a context, we predicted that the response to hearing their sounds would be more similar than the response to the sounds of the *cat* and a *clock*, which did not share a context. Put differently, we predicted that if a specific waveform is involved in the process of contextual reinstatement, spectral power linked with that waveform would be more similar for two sounds that share a context and less similar for two sounds that do not share a context. To test this, we calculated the ICC across sounds for each cluster and each participant (Figure 3d). Higher ICC values would indicate stronger within-context consistency. Results showed higher-than-chance coefficients for cluster 2 (*p*<0.05), and specifically for cluster 2B, reflecting post-sound sigma activity (*p*<0.001). Unlike the previous analysis conducted *within* sounds, this analysis considered similarity *between* sounds and therefore does not reflect trivial sources of correlation, such as acoustic similarity. These results could implicate activity in the sigma band in the process of context reinstatement during sleep.

Finally, we explored a more direct connection between within-context sigma correlations and the aforementioned behavioral effects depicted in Figure 2c. For each participant, we calculated (a) the TMR-induced changes in ESDF values for cued objects and for non-cued objects within cued sets; and (b) the ICC between sounds belonging to the same contextually bound set (for each cluster). We then correlated these measures across participants and found that participants with more similar sigma power within sets also showed less encoding-strength-dependent forgetting for cued objects (*r*=-0.71, *p*<0.01 for cluster 2; *r*=-0.62, *p*<0.01 for cluster 2B; *p*>0.18 for all other clusters; Figure 3e, left). The correlation for non-cued objects, which reflects the contextual effects of cuing on performance, was significant only for the late sigma cluster (*r*=-0.5, *p*<0.05 for cluster 2B, *p*>0.36 for all other clusters; Figure 3e, right).

For this last set of analyses, we used the differences between the ESDF values for cued sets relative to non-cued sets. Our results, indicating a negative correlation between ICC values and encoding-strength-dependent forgetting for cued sets, may therefore just as likely reflect a positive correlation between ICC values and encoding-strength-dependent forgetting for non-cued sets. Indeed, the exact same analysis can be viewed as if it considers the changes in ESDF values for non-cued objects ∉ cued sets relative to the cued objects ∈ cued sets, revealing positive correlations for the sigma clusters (*r*=0.71, *p*<0.01 for cluster 2; *r*=0.62, *p*<0.01 for cluster 2B). Although both these interpretations are equally valid statistically, we argue that the more parsimonious interpretation is that cuing-related ICC values correspond with cuing-related effects, rather than the change in memories that were not biased during sleep. Finally, to complement these analyses, we tested whether overall individual forgetting patterns, regardless of cuing condition, were correlated with ICC values, and there were no significant correlations for any clusters (*p*>0.42). Taken together, results demonstrate that post-cue sigma power is correlated within set and this correlation predicts contextually determined reactivation effects. Our findings therefore support the hypothesis that power in the sigma band reflects contextual reinstatement during sleep.

### Place-specific neural activity during sleep predicts changes in performance

Previous studies operationalized contextual reinstatement during wake using measures of similarity between brain states during encoding and retrieval (e.g.,^7,8^). To test whether place-specific wake-like activity is involved in sleep reactivation, we had participants observe three categories: places, faces, and abstract images (Supplementary Figure 2). The place images included cropped versions of the images subsequently used for the main task (e.g., a movie theater). Using time-series data from all scalp electrodes to train a support-vector machine classifier, we identified clusters of time-points distinguishing places and abstract images (*p*<0.001). The largest cluster spanned between 0.28-0.73 s after image onset during wake (Figure 4a).

**Figure 4:**
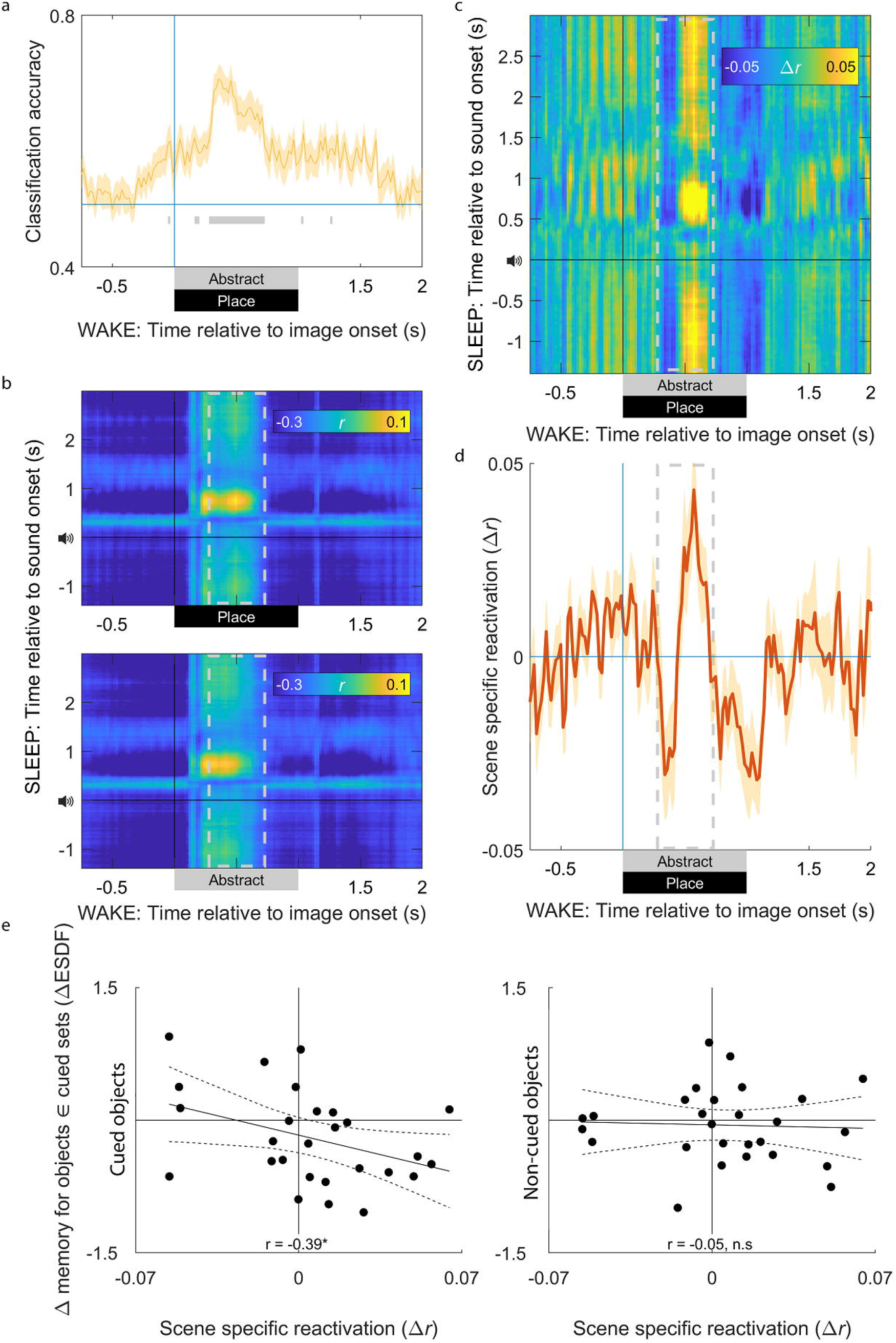
Place-specific neural reactivation predicts changes in performance for cued objects. (a) Classification accuracy for a classifier trained to distinguish images of places and abstract images. Gray lines denote significant classification above chance (*p* < 0.001). Gray dashed boxes in the following panels signify the time period for the largest cluster of continuously significant classification. (b) Correlation patterns between EEG patterns before and after sound presentation during sleep (y-axis) and image presentation during wake (x-axis). Top – correlations for images of places. Bottom – correlations for abstract images. (c) The place-specific correlation patterns, defined as the subtraction between the upper and lower matrices shown in panel b. Note that place-related activity starts before sound onset during sleep and persists throughout most of the sound-locked time-course. (d) Place-specific correlation patterns averaged over the sound-locked time-course during sleep. (e) Correlations between the place-specific patterns during the classifiable time period (gray dashed box in panel d) and changes in encoding-strength-dependent forgetting values across participants. Top – for cued objects ∈ cued sets. Bottom – for non-cued objects ∈ cued sets. P-values reflect the significance level of the Pearson correlation coefficients. Shaded areas and dashed lines signify standard errors of the mean. *-*p*<0.05; n.s-not significant.

We hypothesized that place-related brain networks would play an integral role in offline processing of the scene-heavy contexts used in our task. Therefore, we decided to use the wake, place-specific patterns of activity as a marker for context reactivation during sleep. We next correlated the sleep and wake EEG data from all scalp electrodes to reveal reinstatement of context-related activity. Averaging across wake trials (i.e., around image onset for place images in the functional localizer) and sleep trials (i.e., around sound onset for sleep-related sounds), we calculated the time-point-by-time-point correlation matrix. The result was a time X time matrix of correlation coefficients, with a peak in correlation around 0.5 s after place image onset during wake and 0.75 s after sound onset during sleep (Figure 4b, top). This increase in correlation may reflect genuine neural reactivation of place-related representations. However, it could be that the correlations are not place-specific, but rather reflect similarities between sleep and image-viewing during wake. Indeed, conducting the same correlation analysis between task-related sound presentation data and EEG data following the presentation of an abstract image revealed similar temporal dynamics (Figure 4b, bottom). This similarity suggests that the observed increase in correlation between sleep and wake is driven by wake-like activity occurring during the cuing period that is not necessarily content-specific.

We next tried to tease apart the generalized wake-related correlation patterns and content-specific correlation patterns by considering the differences between the patterns linked with abstract and place images. We defined place-specific reactivation as the subtraction of the place and abstract related matrices (Δr; Figure 4c). The resulting matrix showed a peak in correlation around 0.6 s after place image onset during wake and 0.75 s after sound onset during sleep. Although correlations peaked following sound onset, they persisted at other times as well (including above-baseline correlations before sound onset, which was generally about 6-7 s following prior sound onset). This extended period of place-specific reactivation was unexpected, and seemingly challenges our hypothesis that reactivation during sleep would elicit context-related neural patterns. On the other hand, extended reactivation could stem from features of the TMR design we used; during these critical periods of sleep there were repetitive presentations of task-related sounds, all of which were part of contextually bound sets involving places. We therefore propose that location-related context was reactivated consistently throughout the cuing period, producing reactivation immediately after cues as well as later during inter-sound intervals. Qualitatively similar results were obtained using a classification approach rather than correlation-based analyses (Supplementary Figure 3).

To reveal whether place-specific reactivation predicts changes in performance, we collapsed the place-specific reactivation matrix over the time course of sleep trials (Figure 4d). For the wake data, we focused on periods during which the classifier distinguished between places and abstract images (Figure 4a; gray dashed frame in Figure 4d). Arguably, data from intervals when the two categories are indistinguishable would not be informative in tracking memory-related patterns of activation. The results show periods of time during which Δ*r* was above zero, indicating similarity to place-related wake activity, and periods of time during which Δ*r* is lower than zero. These “dips” could indicate either an increase in similarity to the abstract-image-viewing condition or a negative correlation with the location-image-viewing condition (or both). Future studies should explore these dynamics further, taking into consideration their interactions with sleep-specific waveforms (e.g.,^12^).

We next calculated average correlations during this time period and correlated them with ESDF values for cued objects ∈ cued sets. Results indicated lower ESDF values for cued objects in participants who demonstrated higher wake-sleep place-specific correlations (*r*=-0.39, *p*<0.05; Figure 4e, left). However, reactivation patterns did not significantly correlate with non-cued objects ∈ cued sets (*r*=-0.05, *p*=0.82; Figure 4e, right). These two correlation coefficients were not significantly different from one another *(p*=0.21). Taken together, results support the hypothesis that place-specific neural representations are reinstated during sleep and that this reinstatement predicts encoding-strength-dependent forgetting for cued memories. However, the temporal dynamics of this activity (i.e., its persistence over the intertrial interval) and the lack of a clear correlation with the changes in performance for non-cued objects ∈ cued sets qualify the validity of these results.

## Discussion

In this study, we tested whether the context in which memories are encoded impacts sleep-related consolidation. Participants first developed idiosyncratic stories linking different objects with a physical place, and then encoded on-screen positions for each object. By presenting object-specific sounds during NREM sleep, we selectively biased reactivation towards specific memories, thereby impacting forgetting for these memories. Crucially, this manipulation also impacted retrieval for contextually bound memories that were not cued directly.

If reactivating a memory reinstates its context, carry-over benefits should be determined by the level of contextual overlap between the cued and non-cued memories, with the greatest benefits incurred by the targeted memory itself. Although the impact on cued memories was numerically higher for cued relative to non-cued memories, this difference was not significant. This null effect should be considered with caution, and future studies should more directly consider the hypothesis that the degree of contextual overlap would determine the benefits to contextually linked memories.

For both cued and non-cued memories, observed effects on memory were dependent on initial memory strength. Benefits to memory grew linearly as a function of initial memory strength, with greater benefits for objects that were weakly encoded before sleep and detriments to stronger memories. These data are in line with previous studies that have found that weakly encoded memories seem to be prioritized for reactivation and consolidation. The preferential benefits seem to be a general phenomenon, observed during sleep and resting wake (e.g.,^28^) and impacting both spontaneous reactivation (e.g.,^20^) and targeted reactivation (e.g.,^22,23,29^). However, our data suggest that the beneficial effect of TMR for weaker memories may be mirrored by a detrimental effect for stronger memories. This balance between reactivation-related improvements and detriments in our paradigm raises the possibility that these effects were restricted by some homeostatic process^30^. We did not find a uniform benefit of cuing on memory performance when dependence on encoding strength was not taken into account.

Spectral analyses revealed that power in the sigma band (15-20 Hz) was correlated when sounds were linked to contextually bound objects. Furthermore, these correlations within contextually bound sets predicted changes in retrieval performance: with greater similarity there was less encoding-strength-dependent forgetting for both cued and non-cued objects within a set. The relationship between the physiological and behavioral correlates of contextual reinstatement supports our hypothesis regarding the role of context in consolidation during sleep. Finally, our analyses revealed that neural representations related to places were reinstated during sleep, and this reinstatement predicted changes in retrieval performance for cued objects. However, these results may reflect more general reactivation patterns that are not context-specific and warrant further research.

Taken together, our analyses reveal that context was reinstated during sleep in a manner that had an impact on memory processing. From a broader perspective, these findings fit with the growing literature linking context and memory. The notion of context-dependent memory pertains to improved retrieval in a context similar to the encoding context^31,32^. The natural process of autobiographical retrieval involves the experience of mentally travelling back in time^33,34^, which in itself involves contextual reinstatement ^6^, thereby improving retrieval by increasing the similarity between neurocognitive states at encoding and retrieval.

Despite much theoretical and empirical research on how context bridges memory encoding and retrieval, the question of context’s role in memory consolidation during sleep has been scarcely addressed. Some evidence suggests that sleep serves to strengthen the links between memories and the contexts in which they are encoded^35,36^. Indeed, consolidation over time seems to increase the similarity between neural representations of memories linked with the same context^37^, although the specific role of sleep remains to be explored.

The question of whether contexts are reinstated during sleep is separate from the question of sleep’s effects on context-item binding. Some consolidation theories assume that sleep is a context-less state, and this property of sleep shelters memories by preventing context-related interference^13^. However, to the best of our knowledge, the question of context reinstatement during sleep has not been systematically explored. Recently, we showed that the capacity for reactivation and consolidation during sleep is not limited to a single memory at a given time, suggesting that memories that are tightly and conceptually interlinked can be reactivated simultaneously^19^. Our current results suggest that this capacity for simultaneous reactivation extends beyond tightly interlinked memories (e.g., different memories related to cats) and also applies to contextually interlinked memories (e.g., memories that reside within the same narrative). These findings complement recent studies that have found that reactivating memories during wake had a retroactive beneficial impact on conceptually related memories that were not directly reactivated^38,39^. Taken together, these findings show that consolidation during both wake and sleep involves contextual reinstatement and impacts memories that were not directly cued.

## Limitations of the study

Our design attempted to tease apart two interlinked forms of context – semantic context and temporal context. Results revealed that semantic context, operationalized using idiosyncratic narratives constructed by each participant, impacted sleep-related effects on memory, whereas temporal context, operationalized using a temporally structured block design, did not (Figure 2). However, we acknowledge some design limitations that warrant a nuanced interpretation of this divergence. First, the contextually bound sets, which were used to operationalize semantic context, also shared a temporal context; they not only shared a narrative, but were also learned in the same learning block. Second, temporal context is not impacted exclusively by the mere passage of time. Instead, salient events act as event boundaries, creating abrupt shifts in temporal context^40,41,42^. It could be argued, therefore, that blocks in the position-learning part of the task did not uniformly reflect temporal context, but rather that switches between trials within a block acted as event boundaries. Effectively, this framing would mean that memories in our task never shared temporal context, providing an alternative explanation for the observed null findings.

However, a distinction between the roles of semantic and temporal context during sleep aligns with other experimental findings. In a recent study, we used the same task design to explore the role of context in undisturbed, overnight sleep^43^. By considering the similarities between memory trajectories over a 10-hour delay period within the same contextually bound sets, we found that semantic – but not temporal – context drives performance changes over a period including sleep, but not over a period that did not include sleep. In a separate study, Liu and Ranganath^44^ found the sleep is crucial for binding together memories that are semantically related but learned in different episodes. In contrast, they found that sleep did not impact binding between memories that were semantically unrelated but learned within temporal proximity of each other (see^45^ for computational work supporting this model). These results suggest that there may be a qualitative difference between the roles semantic and temporal contexts play in memory processing during sleep.

A different notable limitation of our design concerns the way sounds were incorporated in our design. The object-congruent sounds were first introduced in the position-learning part of the task (and not earlier, during the story-building part), potentially limiting the sounds’ context-dependent impact when presented during sleep.

Over the last decades, sleep’s active role in memory consolidation has gradually been revealed and acknowledged. Our understanding of how memory representations are reactivated and evolve during sleep is still incomplete. Day-to-day memories are best understood when considering the connections amongst them, yet these connections are not accounted for in our models of memory processing during sleep^46^. Our demonstration of a role for context in sleep consolidation opens the door for further exploration of how memory interconnections impact consolidation during sleep. More generally, this study underscores the notion that memory processing orchestrated by the sleeping brain is as rich and complex as when we are awake.

## Acknowledgments

This work was supported by NIH grants K99-MH122663 and R00-MH122663 and by NSF grant BCS-1921678.

## Author contribution

All authors contributed to the design of this study and helped revise the manuscript. E.S. collected the data. E.S. and J.H. conducted the analyses and E.S. wrote the initial draft of the manuscript.

## STAR Methods

| REAGENT or RESOURCE | SOURCE | IDENTIFIER |
| --- | --- | --- |
| Deposited data |  |  |
| Custom scripts | This paper | https://github.com/cognos-uci/CellReports2023 |
| Software and algorithms |  |  |
| Matlab 2018b | MathWorks Inc, Natick, MA | https://www.mathworks.com/products/matlab.html |
| sleepSMG package | Greer & Saletin | http://sleepsmg.sourceforge.net |
| FieldTrip package | Oostenveld et al., 2011 | https://www.fieldtriptoolbox.org/ |
| Presentation v17.2 | Neurobehavioral Systems, Inc. | https://www.neurobs.com/menu_presentation/menu_download/version_history |
| Other |  |  |
| BOSS image corpus | Brodeur et al., 2014 | https://sites.google.com/site/bosstimuli/ |

### RESOURCE AVAILABILITY

#### Lead contact

Further information and requests for resources should be directed to and will be fulfilled by the lead contact, Eitan Schechtman.

#### Materials availability

This study did not generate new unique reagents.

#### Data and code availability

- The datasets generated during the current study are available from the lead contact upon request.
- Original code has been deposited at GitHub and is publicly available as of the date of publication. DOIs are listed in the key resources table.
- Any additional information required to reanalyze the data reported in this paper is available from the lead contact upon request.

## EXPERIMENTAL MODEL AND SUBJECT DETAILS

We recruited human participants from the local university community who claimed to be able to nap in the afternoon and reported not having a hearing impairment or a history of any neurological or sleep disorders. Participants were asked to go to bed later than usual on the night prior to the study, to wake up earlier than usual on the day of the study, and to avoid caffeine that morning. We assumed that the relevant neurocognitive mechanisms of sleep – which are the core focus of this study – would not be impacted by sleep restriction or participants’ napping patterns. In total, 48 participants were recruited (14 identified as men, 33 identified as women, and one identified as gender queer; average age = 22.6 years). Data from 19 of these participants were excluded from the final analysis (16 who were not exposed to all stimuli during NREM sleep and three with poor recall of which objects were associated with each place, as described below). Although these exclusion criteria were not pre-registered, including these 19 participants was deemed inappropriate because the logic of our manipulation relied on both reactivation during sleep and strong place-object learning. In total, 29 participants were included in the final analysis (8 identified as men, 20 identified as women, and one identified as gender queer; average age = 22.8 years). Based on self-report, participants slept 5.93 hours on average on the night before the study. The Northwestern University Institutional Review Board approved the procedure.

## METHOD DETAILS

### Materials

Visual stimuli were presented on a screen (19201×11080 pixels, P2418HT, Dell Inc., TX). Sounds were delivered over a pair of speakers (AX-210, Dell Inc., TX). Participants’ spoken responses were recorded using a Lavalier clip-on microphone (PoP voice Inc.). Stimulus presentation and participant responses were controlled by Presentation (v17.2, Neurobehavioral Systems, Inc.).

Visual stimuli were used for both for the main task and for the functional localizer task. For the main task, visual stimuli consisted of 76 images of objects and 19 images of places. The object images were square and portrayed either inanimate objects (e.g., a telephone) or animals (e.g., a cat) on a white background. During the spatial task described below, they were each shown at 125⍰×⍰125 pixels (34.4⍰×⍰34.41mm). Most images were taken from the BOSS corpus^47,48^, and some were taken from copyright-free online image databases (e.g., http://www.pixabay.com). Each object image was matched with a distinguishable, congruent sound with a maximal duration of 0.6 s (e.g., a ringing sound; a meow sound). The place images portrayed distinct real-life places (e.g., a movie theater; a desert) and were shown horizontally with a 1:2 aspect ratio. Images were taken from copyright-free online image databases (e.g., http://www.pixabay.com).

Three of the 76 object images and one of 19 place images were used in a pre-task practice block. One additional object image was never displayed, but the sound associated with it was presented during sleep along with a subset of task-relevant sounds, as detailed below. The remaining 18 place images were each associated with four objects to create contextually bound sets. Object images were each assigned a random position on a 2D circular grid (radius – 540 pixels, 148.5 mm). The positions of the 72 objects were set to be at least 50 pixels from the center and the perimeter of the grid and at least 55 pixels from all other object positions. The positions of each set of four objects associated with the same place were at least 425 pixels one from the other. This allowed us to separately estimate errors that stem from confusion between the positions of two objects (swap errors) and errors that stem from imprecise object placement (accuracy errors; see below)^19^.

For the functional localizer, a total of 120 images were used, including 40 images belonging to each of three categories: faces, places, and abstract images. All images were cropped to be square and were presented on-screen at 450 × 450 pixels. The face images were taken from the Psychological Image Collection at Stirling (pics.stir.ac.uk). The place images consisted of the same images used for the main task, cropped, and supplemented by additional images taken from the BOLD5000 database^49^. The abstract images are scrambled place images, created by scrambling the Fourier transforms of place images from the same database.

### Procedure

After consenting to participate in the study, participants were fitted with an electroencephalography (EEG) cap. EEG data was collected continuously throughout all phases of the study (Figure 1a). Since data were collected during the COVID-19 pandemic, participants wore their masks throughout the study, except during the nap portion. After entering the experimental chamber, participants completed a task to measure their response times. This RT task consisted of a red square that shifted between left and right positions at 101Hz and finally stopped at one of the two locations. The participant was required to click the correct mouse button (i.e., left/right) before the square began flickering again. The task ended when the participant responded correctly for 8 out of the last 10 trials. Initially, the square paused for 450 ms, but if the participant failed to reach the criteria within 30 trials, this duration was extended by 50 ms and the task restarted, iterating until the criterion was reached. Then, participants rated their sleepiness level using the Stanford Sleepiness Scale^50^.

Next, participants conducted a functional localizer task (Supplementary Figure 2a). The rationale for including this task was to identify neural patterns that are specific to a category of stimuli. These patterns were then used to identify context reactivation during sleep, as explained below. The task included 150 trials, equally divided among three image categories: places, faces, and scrambled, abstract images. The task included dozens of diverse exemplars from each category to establish that the detected patterns were indeed category specific and not driven by low-level visual features or any one image. Each trial included a 1-s exposure to an image, with an inter-trial interval ranging between 2.5 and 3.5 s. Each category included 40 images, 10 of which were repeated over two consecutive trials. Participants were instructed to left-click the mouse when an image was repeated. The first two participants run did not undergo the functional localizer task, and their data were not used for analyses incorporating data from this task.

Participants next began the main task, which consisted of three parts: story-building, position-learning and test. In the story-building part (Figure 1b), participants were instructed to build idiosyncratic stories, one for each contextually bound set (i.e., images of a place and four objects). Each set was presented together on the screen, and participants had to indicate when they have developed a story for it. Then, they recorded an audio rendition of the story. Finally, they were asked two questions about each object in each story: “Did the object appear throughout the whole story, start to end?” and “was the object in motion (not static) during the story?” These questions were chosen because they were applicable to all objects, yet the answers did not merely concern object attributes but rather required retrieving the constructed story. The answers for both questions with respect to all four objects were recorded before moving on to develop a story for the next contextually bound set.

After story building was completed, participants started the position-learning part (Figure 1c). This part of the task consisted of nine blocks, each including a pair of contextually bound sets and eight objects in total. Each block included interleaving learning trials for the two sets. This design choice was made with the intention of linking the two sets together through a shared temporal context at encoding. Set pairings and block allocation were randomized. In this part of the task, participants had to encode the on-screen positions for the objects on a two-dimensional on-screen circular grid. Once all object positions were learned, as defined below, the next block commenced. Before the first block commenced, participants engaged in a practice block which included four objects that were designated as practice objects.

At the start of each block, participants viewed the images of the two places linked with the two contextually bound sets featured in the block and were given the option to listen to their recordings of the associated stories to refresh their memories. Next, participants were exposed to the to-be-learned position of the eight objects included in the block. Each object was presented in its on-screen position for 4.5 s. Its congruent sound was presented twice, once at trial onset and again at the end of the trial (with sound offset synchronized to image offset). A 1-s inter-trial interval followed.

After being exposed to the true object positions, participants trained on placing the object images in their positions. Each trial included a single object, and trials were presented in a pseudo-random order, such that objects linked to the same story were seldom presented sequentially. At the start of each trial, an object-specific contextual question was presented. These questions were the same ones presented in the story-building part. Participants had to get each question correct to proceed with the trial (responses that were congruent with the answers recorded previously were deemed correct). Answering incorrectly terminated the trial (Figure 1c, bottom). The purpose of presenting these questions was to repeatedly reinstate the encoding context during the position-learning part of the task. Overall, 88.92%±0.9% (mean ± SEM) of questions were answered correctly during training. Next, the object was presented at a random position on the grid, along with its associated sound. Participants used the computer mouse to attempt to drag the object to the position where it was initially seen on the grid. Trials with a Euclidean error of less than 100 pixels relative to the true position were considered correct. After correctly placing an object near its true position twice in a row, it was considered as learned and was dropped out from the block. On average, each object was presented in 3.52±0.15 trials (mean ± SEM). Both correct and incorrect trials included feedback: the true object position was presented for 2 s along with the user-selected position. The sound was then presented again, co-terminating with the feedback display. A 1-s inter-trial interval followed.

The last part of the main task including a test on object positions. In each trial, participants had to drag one of the objects to its true position. All 72 objects were presented in a pseudorandom order, preceded by the three practice objects. Objects were each presented in a random position on the grid, accompanied by their sounds. No context-related questions were presented, nor was feedback given. A 1-s inter-trial interval was used.

Following the test, participants’ pre-sleep error rates were calculated (i.e., the Euclidean distance between the chosen and true on-screen positions). Out of a total of 72 objects, 12 were designated to be cued during sleep. These objects were selected in a manner that obeyed the following logic: Six of the nine blocks included cued objects. Each of these blocks consisted of two contextually bound sets: one including cued objects; the other not. The six contextually bound sets which included cued objects each included two cued objects and two non-cued objects. Out of the nine blocks, the remaining three blocks did not include any cued objects. The condition designated to each object, set, and block were determined using an algorithm that minimized variability between the average error rates among conditions. For each participant, the algorithm considered 200,000 random allocations of objects to conditions, each obeying the logic outlined above. The random allocation for which the average positioning error was lowest across the following four conditions was used: (1) cued object ∈ cued set; (2) non-cued object ∈ cued set; (3) (non-cued object ∉ cued set) ∈ cued block; (4) non-cued object ∉ cued block. In addition to the 12 object-related sounds designated for cuing, another sound which was not used during the wake portions of the task was presented during sleep as a control sound.

Immediately after ending the test, participants were permitted to nap for 90 minutes with the lights out on a foldable futon in the same experimental room. Throughout their nap, white noise was presented (∼47 dB). Sleep was monitored online by an experimenter skilled at sleep staging. Upon detection of stage N3 of NREM sleep, sounds were unobtrusively presented in the experimental room (<53 dB). EEG data were monitored continuously, and sound presentation was terminated immediately upon signs of arousal or transition to REM sleep. The inter-stimulus interval (i.e., offset-to-onset) was randomly set to either 6, 6.5, or 7 s. If a participant did not reach NREM stage N3 after 45 minutes, sounds were presented in either stage N2 or N3 throughout the remainder of the nap. Out of the 48 participants, 16 participants were not exposed to all 13 sounds at least once during NREM (they were exposed to 4.75±0.83 sounds on average throughout all sleep stages). These participants either never reached stable NREM sleep or were easily arousable throughout the nap. Since sound presentation served as the main manipulation in this study, these participants were excluded from all analyses. Although the excluded and included participants may differ in terms of sleep quality, potentially introducing a selection bias, it is highly unlikely that these groups would differ in the relevant mechanisms of memory processing during sleep.

After the nap, participants were required to wait at least 5 minutes before resuming the task. Before completing the post-sleep memory tests, participants engaged in the response-time task described above. To rule out the effects of sleep inertia on performance^51^, participants were required to meet a response-time criterion that was set based on their pre-sleep performance on the same task. Additionally, participants rated their sleepiness level once more. The result for both tasks are detailed in Supplementary Table 2. Next, they started the post-nap test, which was identical to the pre-nap test. Then, participants completed a self-paced recall test, in which they had to type in, for each picture of a place, which objects were linked with it. This part of the task was used as a manipulation check, since the expected effects of TMR critically depended on a strong, over-trained link between objects, stories, and places. Three participants who failed to recall at least 75% of the objects were excluded from analysis. Finally, participants were asked if they heard sounds presented during the nap. Out of the 29 participants used for analyses, 7 reported hearing task-related sounds. These 7 participants then underwent a task in which they were required to indicate which sounds they remember hearing during sleep. Their responses indicated that they were not significantly different than chance at identifying which sounds were presented (*p* = 0.8, Sign Rank Test). Participants were then allowed to clean up, after which they were dismissed.

### Electrophysiological data collection and preprocessing

EEG was recorded using Ag/AgCl active electrodes (Biosemi ActiveTwo, Amsterdam). In addition to the 64 electrodes at 10-20 system scalp locations, contacts were placed on the mastoids, next to the eyes, and on the chin. Recordings were made at a sampling rate of 512 Hz. Analyses were conducted using the FieldTrip^52^ and sleepSMG (http://sleepsmg.sourceforge.net) packages for Matlab 2018b (MathWorks Inc, Natick, MA). EEG channels were re-referenced offline to averaged mastoids and filtered using a two-way least-squares FIR highpass filter with a cutoff of 0.4 Hz. Additionally, a notch filter was used to remove noise at 60 Hz. Noisy channels were replaced with interpolated data from neighboring electrodes using the spherical interpolation method in FieldTrip, and noisy segments were detected manually and removed from further analyses. For the data collected during wake, ICA was used to detect and remove artifacts associated with eye blinks and horizontal eye movements.

### QUANTIFICATION AND STATISTICAL ANALYSIS

#### Sleep staging

Sleep staging (i.e., determining the stage of sleep for each 30-s epoch) was based on the guidelines published by the American Academy of Sleep Medicine^53^ and conducted by two independent raters, both of whom were not privy to when sounds were presented. Any discrepancies were subsequently reconciled by one of the two raters. Supplementary Table 1 shows the amount of time spent in each stage of sleep and number and percentage of cues presented in each stage.

#### Statistical analyses of behavioral data

For each trial in the tests conducted before and after sleep, the error was measured in pixels as the Euclidean distance between the true object’s position and the position indicated by the participant. In designing this experiment, we anticipated that the positions of contextually linked objects, which were learned within temporal proximity, would be confusable. Therefore, to disentangle any gross miscategorization errors from nuanced positioning errors (and focus on the latter, which were central to the design of this study), our algorithm positioned the four objects within a set at a substantial distance from one another (425 pixels). This value was over three times larger than the median error across participants and conditions (82.8 pixels), allowing us to categorically flag objects as “swapped” if they were placed closer to the position of another object belonging to the same contextually bound set. These gross errors, stemming from either pure guesses or confusion between objects, were omitted from further analysis^19^. In total, 20.07%±1.81% of objects were swapped before sleep and 19.88%±1.99% after sleep. The change in swap rates over sleep was not modulated by condition *(F*(3, 2084)=1.41, *p*=0.24; *Change_in_swap_rate ∼ 1 + Condition + (1 + Condition* | *Participant))*. For all the remaining trials, repeated-measures ANOVAs and mixed linear models were used. All repeated-measures ANOVA use the Lower-Bound Estimate to correct for violation of the assumption of sphericity. Results were comparable to those obtained with other methods (i.e., Greenhouse-Geisser correction; Huynh-Feldt correction) or without any correction. The following mixed linear model was used for comparing pre-sleep accuracy errors between conditions across participants (*fitglme* function in Matlab):

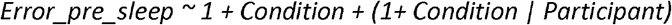

“Participant” is a categorical variable, denoting the participant number of each individual participant. Two different analyses were run, one focusing only on semantic context and one considering semantic and temporal context separately. For the former, “Condition” was a categorical variable with three possible values: (1) cued object ∈ cued set; (2) non-cued object ∈ cued set; (3) non-cued object ∉ cued set (Figure 1d). For the latter, “Condition” was a categorical variable with four possible values: (1) cued object ∈ cued set; (2) non-cued object ∈ cued set; (3) (non-cued object ∉ cued set) ∈ cued block; (4) non-cued object ∉ cued block (Figure 2d). An ANOVA was used to report the statistical significance of the components of the model, and dummy variables were used for comparisons between conditions (producing the p-values reported in the paper and presented in Figure 2).

To consider the changes over sleep, error rates were Z-scored within participants and used to evaluate the effects of cuing during sleep across participants using the following mixed linear model:

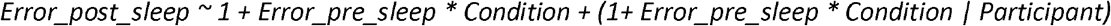

This analysis was motivated by recent studies suggesting that targeted memory reactivation in these experimental paradigms (and likely spontaneous sleep reactivation as well) selectively improves memory for weakly encoded memories^22,23,29^. By using normalized data and adding the pre-sleep error as a predictor, we were able to separately evaluate two effects: a uniform cuing benefit, which would manifest as a main effect of Condition (a different in intersect in Figure 2b); and cuing effects that rely on pre-sleep error levels, which would manifest as an interaction effect between Condition and Error_pre_sleep (a different slope in Figure 2b, 2c; encoding-strength-dependent forgetting). A similar analysis was run on non-Z-scored data (Supplementary Figure 1).

In order to quantify the extent of the cuing effect on cued and non-cued objects within a contextually bound set for each participant (Figure 3e, Figure 4e), a similar model was run for each participant, omitting the random effects.

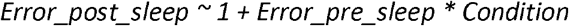

The obtained coefficients for each condition [i.e., condition-specific encoding-strength-dependent forgetting values relative to the value for the (non-cued object ∉ cued set) group] were then used to calculate correlations between behavior and physiology across participants, as outlined below.

#### Spectral analysis

Spectral analyses were run on data collected during sleep from electrode Cz and limited to epochs for task-related sounds presented during NREM sleep. We decided to focus our analyses on electrode Cz because it is sensitive to two notable memory-related waveforms: sleep spindles and slow-oscillations^19^. Trials were segmented around sound onset (1.25 s before to 4.75 s after). For each trial, we first subtracted its overall mean and then calculated a spectrogram between 0.25 Hz and 25 Hz in 0.25 Hz intervals, using 0.5-s time windows with 87.5% overlap. For each participant, the average baseline (i.e., *t* < 0 s) activity was calculated per frequency band, and each trial’s spectrogram was converted to percent change by subtracting and dividing the activity during baseline for each frequency band. These trial-specific spectrograms were used to extract power in specific time-frequency clusters on the single trial level, as detailed below.

To identify significant clusters of sound-related activity, we first averaged the trial-specific spectrograms within participants. Then, each point in the time-frequency representation was compared to zero across participants, with an alpha level of 0.01 (corrected for the number of data-points using a Bonferroni correction). For example, all data points across participants representing t = 1 s, F = 10 Hz were submitted to a t-test with the null hypothesis that the power is zero. This comparison indicated which points in time-frequency space show activity significantly higher than baseline. The results, shown in Figure 3b, indicated there were several clusters of points that were temporally and spectrally contiguous and separate one from the other. The two largest clusters were identified by our algorithm and used for further analyses. The higher-frequency cluster, reflecting activity in the sigma range (see Figure 3b), may encapsulate sleep spindle activity which commonly commences approximately 1 s after sound onset (e.g.,^11,19^). We therefore considered both the full cluster as well as the two putatively separate components, in our analyses.

#### Intraclass correlations of spectral power

For each trial, we extracted and summated the values confined by each cluster from the spectrogram, resulting in a single scalar value per trial and cluster. We hypothesized that trials involving the same sounds within the same participants would have correlated power in certain clusters. To test this, we used intraclass correlation^54^. This metric, ICC, is symmetrical (i.e., whereas *inter*-class correlations predict Y from X, *intra-*class correlations predict how clustered together different values of X are) and can be used to calculate the correlation between more than two values. Unlike interclass correlations, ICC values so not have a minimal value (i.e., can go lower than -1), and are negatively biased (see Supplementary Figure 4). We calculated ICCs for each participant and cluster, and then ran a permutation test with mixed labels (*n* = 10,000) for each participant. Finally, we conducted a paired t-test between the true ICC and the average ICC value calculated using the permutation test. This analysis resulted in the p-values that were presented in Figure 3 and in the main text. To ensure that we had sufficient data and to avoid biases due to a small number of trials, ICC analyses on spectral data were limited to participants who had at least five presentations for each sound (*N* = 16). For these participants, data for all artifact free sound-locked trials was used (average number of trials = 7.5).

Next, we hypothesized that contextually bound memories would elicit correlated spectral activity. We leveraged the fact that for each contextually bound set, two sounds were presented. We averaged the power per sound and per cluster, producing two values for each contextually bound set within participants. The ICC was then calculated within set for each cluster. Like before, we used a permutation test (*n* = 10,000) and paired t-tests to consider evidence in support of our hypothesis. Finally, the true ICC values obtained for each participant and cluster were then correlated with the encoding-strength-dependent forgetting values calculated per participant. The p-values derived from this analysis are presented in Figure 3 and in the main text.

#### Classification analyses

Using the data collected in the functional localizer task, a classifier was trained for each participant to distinguish faces, places, and abstract images (Supplementary Figure 2b). For each image, trials that were not contaminated with artifacts were segmented between 1.5 s before and 3.5 s after image onset. Classification was calculated for each time point independently using time-series data from the 64 scalp electrodes as features. Data were smoothed using a 51.2-ms rectangular smoothing window. These data were used to train a support vector machine (SVM) classifier and tested using 5-fold cross-validation protocol^55^. This procedure was repeated for 20 iterations and averaged across iterations. To calculate the significance level, clusters of contiguous time points that were significantly higher than chance (p < 0.001) were identified, and each time point’s t-values were summed together to produce a cluster-level t mass. Then, a permutation test was initiated by reconducting the classification analysis and identifying significant clusters using shuffled labels. Significance for each true cluster-level t mass was evaluated relative to the random distribution of clusters generated based on 100,000 permutations, with an alpha of 0.001.

#### Sleep-wake electrophysiological pattern correlations

Having established a link between semantic contexts and images of locations during wake, we tested whether the reinstatement of context during sleep would result in an increase in the correlation between neural activity during sleep (reactivation) and wake (place-image viewing). As a baseline condition, we initially considered using the EEG activity related to face-image viewing, but opted against this option in the analysis stage, following the realization that we do not have an *a priori* assumption regarding the involvement of face representations as part of the semantic context (e.g., some contexts may involve people whereas others may not, adding noise to this analysis). Instead, we decided to use the abstract-image-related wake activity as a baseline condition.

Using the Functional Localizer data and the data from the sleep phase, we correlated (1) place-image-related and abstract-image-related wake EEG patterns with (2) patterns observed around the onset of sounds during sleep. First, time-series data were segmented around both the wake and sleep trial onsets, starting 1.5 s before and ending 3.5 s after stimulus onset. Trials containing artifacts were omitted. Only sleep trials that included task-related sounds were considered. Data was smoothed using a 51.2-ms smoothing window. Data from the 64 scalp electrodes at each time point during wake were correlated with data from the same electrodes at each time point during sleep, resulting in a time X time correlation matrix showing wake-sleep correlations across time points. Two matrices were created, one correlating place-image-related wake EEG patterns with sleep-related EEG patterns, and one correlating abstract-image-related wake EEG patterns with sleep-related EEG patterns. The subtraction between the two was used to assess evidence for place-specific activation patterns. The difference matrix was collapsed over the sleep-time axis to create a vector of correlation coefficients over the course of the wake trial, and the time period during which wake classification was significantly above chance was extracted. These values, across participants, were then correlated with the encoding-strength-dependent forgetting values calculated per participant. The p-values derived from this analysis are presented in Figure 3 and in the main text.

