## Supplementary Materials for "Memory consolidation during sleep involves context reinstatement in humans"

**Supplementary Table 1: Time asleep and cuing statistics in each sleep stage**

| Sleep stage | Wake | Stage N1 | Stage N2 | Stage N3 | REM |
|---|---|---|---|---|---|
| **Minutes in stage (mean ± SEM)** | 20.67±2 | 14.91±1.4 | 33.55±2.5 | 17.84±2.3 | 5.86±1.8 |
| **% Time in stage (mean ± SEM)** | 22.41±2.2 | 16.08±1.6 | 36.11±2.7 | 19.29±2.5 | 6.11±1.8 |
| **# Cues in stage (mean ± SEM)** | 0.41±0.1 | 0.62±0.3 | 19.03±4.4 | 57.03±9.4 | 0.28±0.24 |

<u>Supplementary Table 2:</u> **Results for the Stanford Sleepiness Scale before and after sleep and the response-time task**

| | **Stanford Sleepiness Scale** (mean ± SEM) | **Number of trials until criterion** (mean ± SEM) | **Response time in ms** (mean ± SEM) |
|---|---|---|---|
| **Pre-sleep** | 2.83±0.2 | 15.14±1.34 | 384.36±4.19 |
| **Post-sleep** | 2.9±0.18 | 12.79±1.03 | 383.25±3.56 |
| **Pre vs Post (two-way)** | $t(28) = 0.31$, $p = 0.76$ | $t(28) = 1.29$, $p = 0.21$ | $t(28) = 0.26$, $p = 0.8$ |

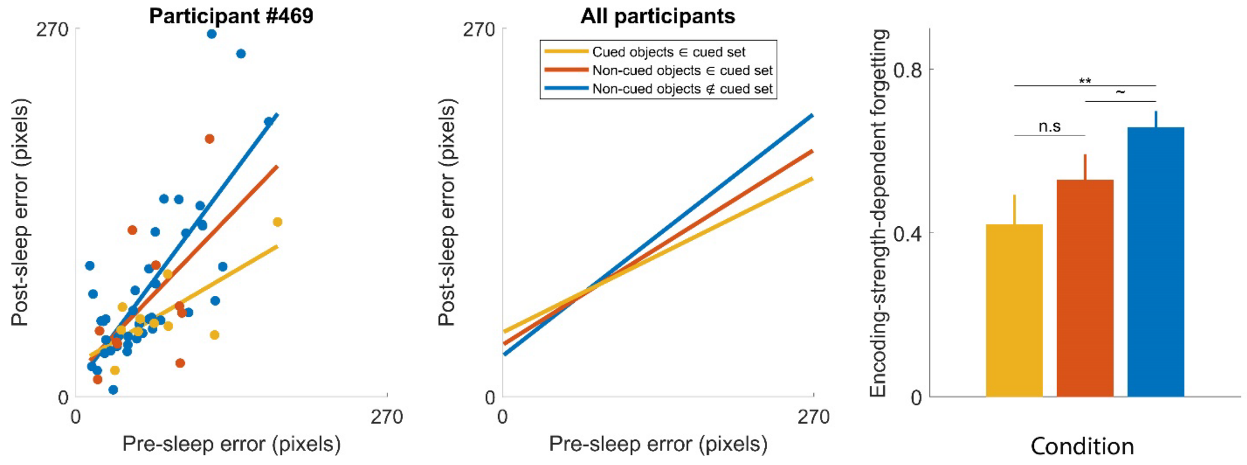


**<u>Supplementary Figure 1:</u> Forgetting curves across all participants considering un-normalized accuracy errors, related to Figure 2.** The left, center and right panels correspond to panels a, b and c in Figure 2, respectively. Unlike in Figure 2, results here were obtained using data that were not Z-scored. Error bars signify standard errors of the mean. ** - $p < 0.01$; * - $p < 0.05$; ~ - $p < 0.1$; n.s - $p > 0.1$.

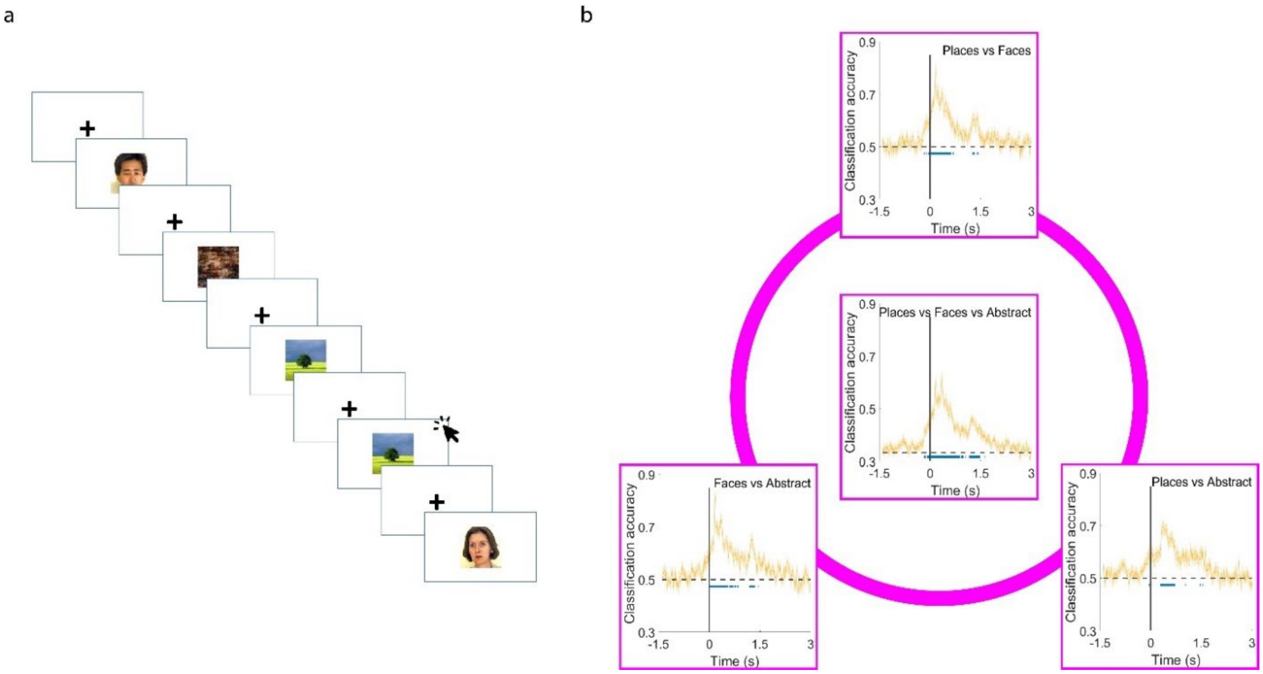


**<u>Supplementary Figure 2:</u> Functional localizer task and classification during wake, related to STAR Methods.** (a) Participants were presented with images and required to indicate when two successive images were identical (i.e., a 1-back task). (b) EEG data collected during the task were used to train an SVM classifier to distinguish between image categories. For all four comparisons, classification was significantly higher than chance. Yellow shades indicate the standard errors of measurement between participants. Blue lines indicate classification significantly above chance ($p$ < 0.001).

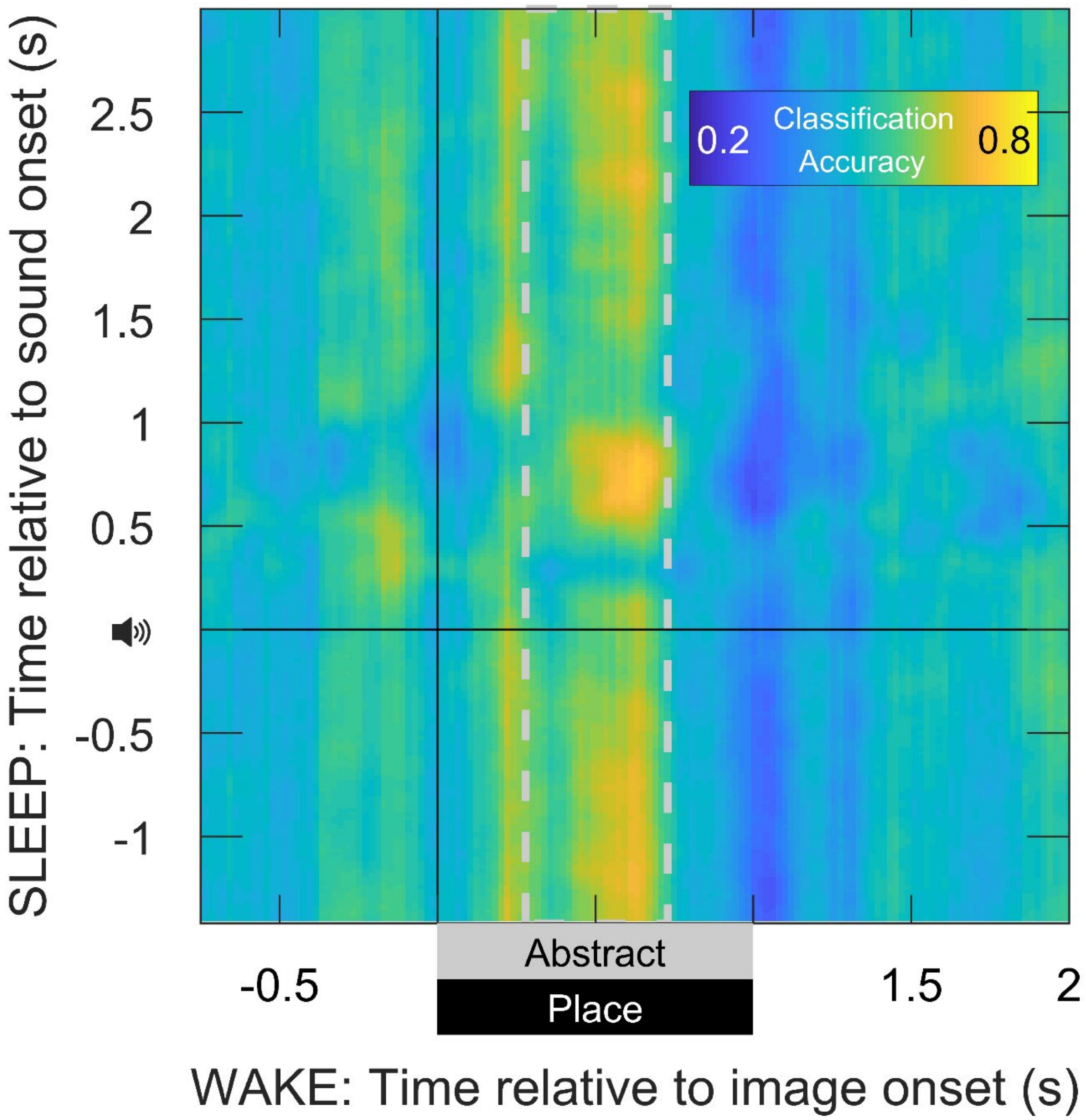


**<u>Supplementary Figure 3:</u> Place-specific neural reactivation as detected using a classification approach, related to Figure 4.** To complement the analyses presented in Figure 4, we subjected the same data to a classification approach rather than a correlational one. We first trained SVM classifiers using all the data obtained from the Localizer task (see Methods). A classifier distinguishing places and abstract images was calculated for each time point around image onset. Then, these classifiers were used to classify the EEG data collected around sound onset during sleep. The figure depicts the probability of each sound-locked time point to be classified as "place" based on a specific classifier. The color scale reflects the probability for each time point to be designated as reflecting place-related activity (chance level = 0.5). Similar to the results presented in Figure 4c, we observed continuous evidence for reactivation around the onset of sounds during sleep (but note that these analyses are not independent of one another).

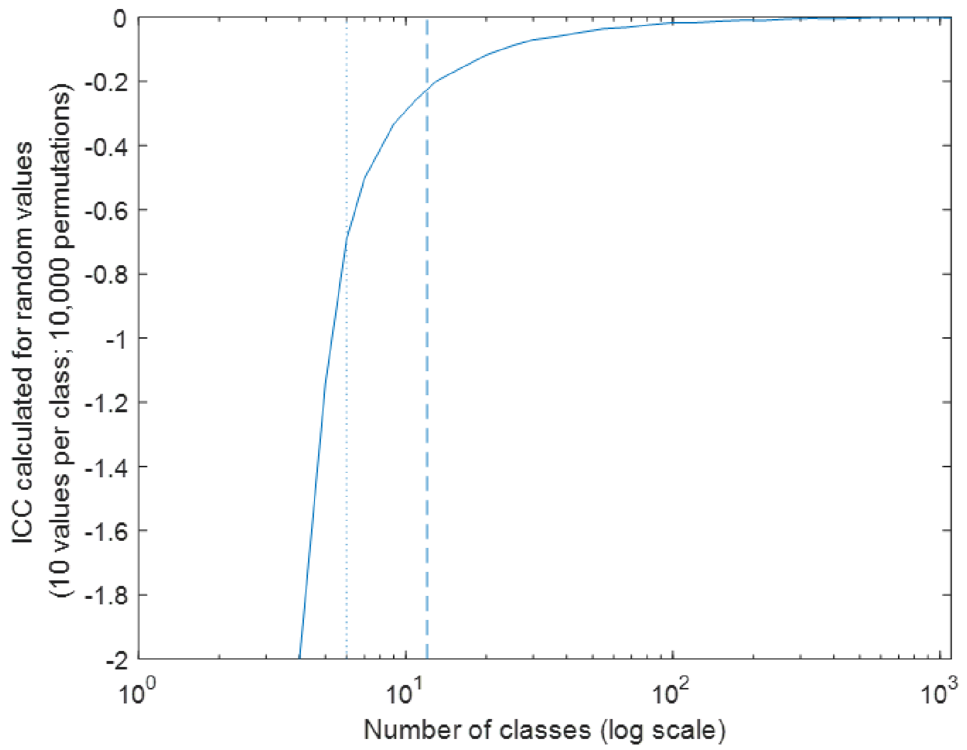


**<u>Supplementary Figure 4:</u> Simulation results demonstrating the negative bias for ICC (1-k) as a function of the number of classes, related to STAR Methods.** In our analysis, we used the (1-k) formulation of ICC, which can be seen as a way to analyze variance between vs. within groups. By definition, positive ICCs indicate that most of the variability is explained by differences between groups (e.g., power in the sigma range after a sound was similar across repetitions of the same sound, but different across sounds). Negative values suggest that most of the variability is explained by differences within group (e.g., the differences between repetitions are greater than those between sounds). However, the ICC is negatively biased in some of its formulations (e.g., Atenafu et al., 2012), producing negative value for shuffled data, as can be seen in Figure 3. To demonstrate this, we ran simulations showing that the negative bias is dependent on the number of classes used. The dotted and dashed lines reflect the values 6 and 12, respectively, which are the values used in the analyses in Figure 3. For these simulations, we used random values chosen from a uniform distribution between 0 and 1, and arranged them randomly into classes, with 10 values in each class. This was repeated 10,000 times for each number of classes and averaged across permutations. The results show that fewer classes result in more negative ICCs for this random dataset. The values we received for datasets of 12 and 6 classes are shown in dashed and dotted lines, respectively. These values closely match the values obtained for our shuffled data as presented in Figure 3c (12 classes) and Figure 3d (6 classes). Importantly, these biases do not impact our results, as we never compared any ICC value with 0. Such comparison would be severely impacted by these biases. Instead, our analyses compared a dataset with the same values rearranged in permutations tests. The real and shuffled datasets should be similarly impacted by the biases demonstrated by this simulation, making the comparison between the two results a valid measure of how correlated values are to one another.
